# Anatomical Abnormalities Suggest a Compensatory Role of the Cerebellum in Early Parkinson’s Disease

**DOI:** 10.1101/2024.08.30.610437

**Authors:** Juyoung Jenna Yun, Anastasia Gailly de Taurines, Yen F Tai, Shlomi Haar

## Abstract

Brain atrophy is detected in early Parkinson’s disease (PD) and accelerates over the first few years post-diagnosis. This was captured by multiple cross-sectional studies and a few longitudinal studies in early PD. Yet only a longitudinal study with a control group can capture accelerated atrophy in early PD and differentiate it from healthy ageing. Accordingly, we performed a multicohort longitudinal analysis between PD and healthy ageing, examining subcortical regions implicated in PD pathology, including the basal ganglia, thalamus, corpus callosum (CC), and cerebellum. Longitudinal volumetric analysis was performed on 56 early PD patients and 53 matched controls, with scans collected 2-3 years apart. At baseline, the PD group showed a greater volume in the pallidum, thalamus, and cerebellar white matter (WM), suggesting potential compensatory mechanisms in prodromal and early PD. After 2-3 years, accelerated atrophy in PD was observed in the putamen and cerebellar WM. Interestingly, healthy controls – but not PD patients – demonstrated a significant decline in Total Intracranial Volume (TIV), and atrophy in the thalamus and mid-CC. Between-group analysis revealed more severe atrophy in the right striatum and cerebellar WM in PD, and in the mid-posterior CC in controls. Using CEREbellum Segmentation (CERES) for lobule segmentation on the longitudinal PD cohort, we found a significant decline in the WM of non-motor regions in the cerebellum, specifically Crus I and lobule IX. Our results highlight an initial increase in cerebellar WM volume during prodromal PD, followed by significant degeneration over the first few years post-diagnosis.

## Introduction

Extensive research indicates that Parkinson’s disease (PD) involves the degeneration of the basal ganglia (BG), particularly the dopaminergic neurons within the Substantia Nigra (SN), leading to the hallmark motor deficits observed in the disease (Marsden, 1990). The neurodegenerative process in PD is characterised by a critical loss of striatal volume, with atrophy in the caudate and putamen, which are integral to the cortico-basal ganglia-thalamo-cortical loop involved in motor control (Pitcher *et al*., 2012). The atrophy and disrupted neuronal activity within these regions undermine the equilibrium between the direct and indirect motor pathways, resulting in the movement impairments characteristic of PD (Szewczyk-Krolikowski *et al*., 2014).

Notably, by the time motor symptoms become apparent in PD patients, over 50% of the dopaminergic neurons in the SN have been lost (Marsden, 1990). This neuronal loss correlates with the deterioration of the basal ganglia’s functionality, as evidenced by reduced resting functional connectivity and the subsequent decline in cognitive abilities as the disease progresses (Szewczyk-Krolikowski et al., 2014; Lopes et al., 2016). Furthermore, the striatum’s volume has been positively associated with ‘motor reserve’, a concept describing the capacity to counteract motor decline by adapting relevant neural networks, highlighting its significance in the early stages of PD (Jeong et al., 2022; Hoenig et al., 2023).

In contrast to the decades’ long focus on the BG itself, recent studies started to shed light on the potential compensatory role of the cortex (Johansson *et al*., 2024) and the cerebellum (Simioni, Dagher and Fellows, 2016; Li, Le and Jankovic, 2023), and their connections with the BG in the cerebello-thalamo-cortical loop (Abulikemu, Tai and Haar, 2023). Recent work, including a cross-sectional study by Kerestes et al. (2023), identified abnormal cerebellar volumes in PD patients compared to a control group. This study underlined the potential link between early-stage PD and dysregulation in the anterior cerebellum and alterations in the posterior non-motor lobes in the later stages of the disease.

Therefore, our study seeks to investigate the volumetric changes within these key brain structures in early PD, over the first 2-3 years post-diagnosis, utilising T1-weighted MRI data. This longitudinal perspective is crucial to identify compensatory mechanisms in prodromal and early PD and to reconcile the diverse findings of previous studies. By analysing both the grey and white matter alterations over time, we aim to provide a clearer picture of PD’s progression, which could contribute to a better understanding of PD.

## 2. Methods

### 2.1 Participants

In this study, we analysed MRI scans for a cohort of people living with PD from the Parkinson’s Progression Markers Initiative (PPMI; see https://www.ppmi-info.org), and the cohort of healthy aging from the Open Access Series of Imaging Series 3 (OASIS-3; see https://sites.wustl.edu/oasisbrains/home/oasis-3/). As the PPMI dataset did not provide a sufficient age-and-sex-matched healthy controls, we obtained these controls from the OASIS-3 dataset. From the PPMI, we acquired clinical and MRI data of PD patients, who have had at least two scans 2-3 years apart, with motor symptoms such as tremor or bradykinesia, scanned within two years from diagnosis, and medication-naïve prior to baseline scan. The patients were all medicated by the time the second scans were acquired. Exclusions included any significant cognitive impairments or comorbidities that could affect the MRI scan. Two cognitive rating scores were used when selecting participants: the United Parkinson’s Disease Rating Scale (UPDRS) and the Montreal Cognitive Assessment (MoCA). Here we included only participants who had an initial score for the UDPRS-part I cognitive impairment of 0, no neuropathological conditions like Alzheimer’s Disease or Chronic Traumatic Encephalopathy, and a MoCA score above 26. This is to ensure participants exhibited no significant cognitive impairment which may confound our results. Individuals with a score below 26 have previously been found to display difficulties completing cognitive and motor tasks, despite intact motor function (Nazem *et al*., 2009). In addition to controlling for the confounding factors, we also minimised motion artefacts by checking for the MRI quality documentation provided by the PPMI dataset. Here, we disregarded participants with poor or “nan” scores. This was cross-analysed through manual inspection of the MRI scans.

The control cohort was derived from the OASIS-3 dataset, which comprised cognitively normal adults aged between 42 to 95 years, without a biological parental history of Alzheimer’s disease, and who had undergone regular cognitive assessments over 15 years. Any indications of cognitive or functional decline reported by cohabitants or comorbidities found in MRI scans resulted in exclusion from our longitudinal dataset.

After the initial selection, demographic outliers were eliminated if they exceeded 3 standard deviations (SD) from the mean. Following screening, our cohort consisted of 56 PD patients (35 men and 21 women, aged 62.44 ±9.62 years, mean ± SD) and 53 controls (32 men and 21 women, aged 64.77 ±7.47 years, mean ± SD). There was no significant difference in age and gender between the two groups (Table 1). The scan year difference within the control group (2.54 ±0.68 years, mean ± SD) was significantly greater than in the PD group (2.09 ±0.29 years, mean ± SD). Table 1 summarises the demographic and clinical characteristics of the participants.

**Table 1.** Demographic and Clinical Data of the PD patients and healthy controls. Age, Disease duration, Scan Difference, UPDRS III, IV, and MoCA scores are calculated in mean ± SD, whereas Hoehn and Yahr (H-Y) PD functional disability scale and UPDRS-I scores are calculated in median (range).

|  | Control (n=53) | PD (n=56) | t/x<sup>2</sup> | p-value |
| --- | --- | --- | --- | --- |
| **Gender, M/F** | 32/21 | 35/21 | 0.00 | 0.97 |
| **Age, year** | 64.77 ±7.47 | 62.44 ±9.62 | 1.40 | 0.16 |
| **Disease duration, years** | - | 2.28 ±0.80 | - | - |
| **Scan Difference, years** | 2.54 ±0.68 | 2.09 ±0.29 | 4.55 | 1.47e-05 |
| **H-Y stage, Baseline** | - | 1 (1-2) | - | - |
| **H-Y stage, Post-2/3 years** | - | 2 (1-3) | - | - |
| **UPDRS-I, Cognitive Score** | - | 0 (0-1) | - | - |
| **UPDRS-III, Total Score (Baseline)** | - | 21.6 ±8.94 | - | - |
| **UPDRS-III, Total Score (Post-2/3 years)** | - | 24.8 ±11.9 | - | - |
| **UPDRS-IV, Motor Complications** | - | 0.55 ±1.93 | - | - |
| **MoCA** | - | 27.3 ±1.93 | - | - |

### 2.3 Imaging Acquisition

T1 MRIs were selected for matching dimensions and voxel sizes, ensuring a well-regulated comparison. The field strength, scanner manufacturer, echo time (TE) and repetition time (TR) were all matched between the two datasets (Table S1). While there were differences in TE and TR times, those were not statistically significant (p = 0.27 for TE and p = 0.054 for TR). For all scans in both cohorts, the T_1_-weighted MRI data consisted of 176 slices with a matrix size of 256×256 and voxel sizes of 1.0×1.0×1.0 mm in plane resolution. Participants who did not meet these parameters were excluded from the study.

### 2.4 T_1_–weighted MRI Data Preprocessing

MRI data were preprocessed with the Freesurfer software package 7.3.2 (Fischl, 2012). This includes the removal of non-brain tissue, and segmentation of subcortical grey and white matter based on image intensity. Individual brains were registered to an atlas parcellating the subcortical regions of interest (ROIs) using the Destrieux anatomical atlas (Destrieux et al., 2010). Based on our interest in the Cortico-basal ganglia-thalamo-cortical (CBTC) and the cerebello-thalamo-cortical (CTC) circuits, the subcortical ROIs analysed were: Caudate, Putamen, Pallidum, Thalamus, and Cerebellar Grey and White Matter (Fig. 1). Other white matter regions, such as cerebral white matter and corpus callosum (CC), were included due to previous findings of their involvement in PD (Taylor *et al*., 2018).

**Figure 1.**
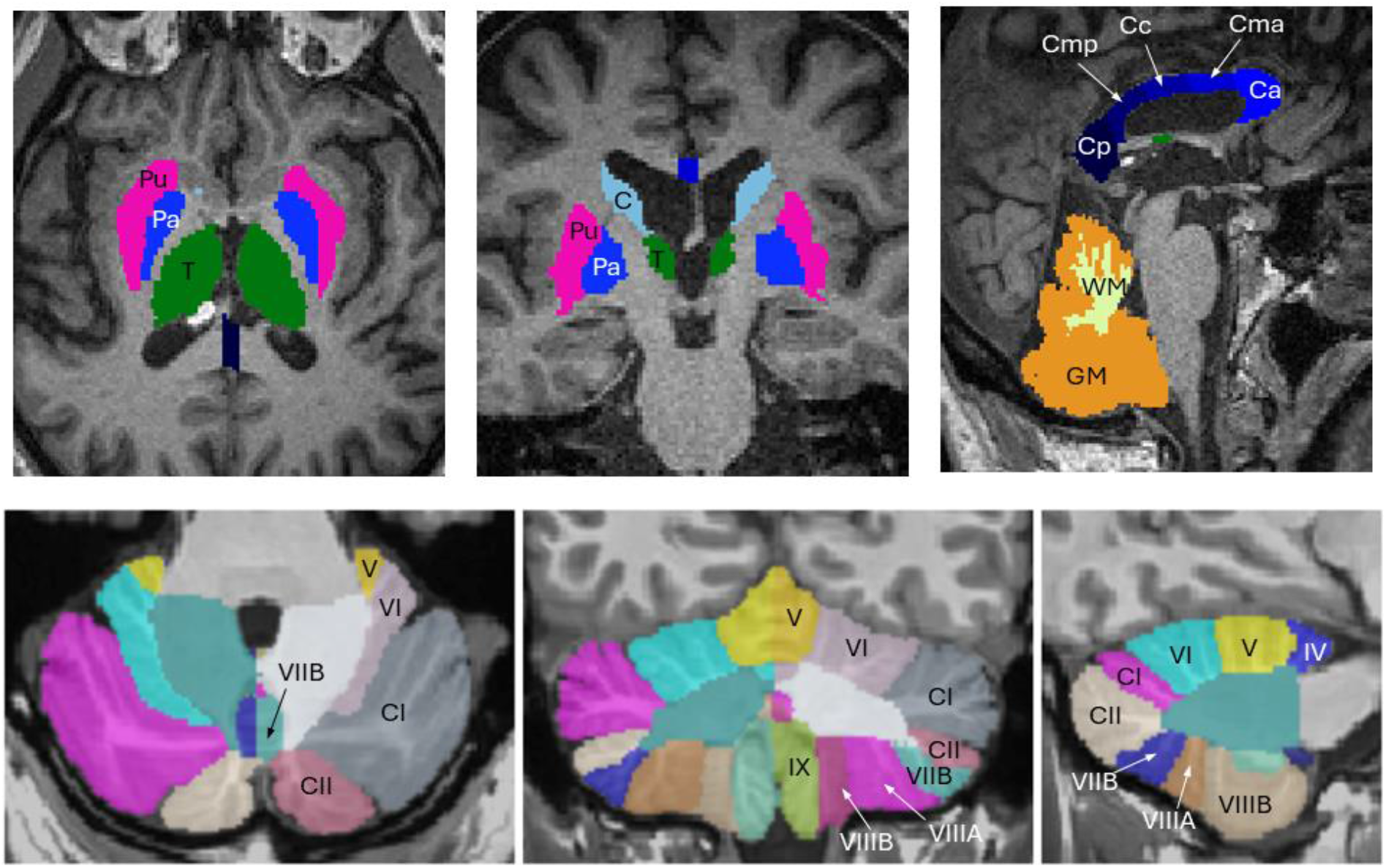
Anatomical regions highlighted on a T1-weighted MRI scan. The upper panel includes the subcortical ROIs from the Freesurfer Destrieux anatomical atlas: **T** = thalamus, **Pu** = putamen, **Pa** = pallidum, **C** = caudate, **WM** = cerebellum white matter, **GM** = cerebellum grey matter, **Cp** = posterior CC, **Cmp** = mid-posterior CC, **Cc** = central CC, **Cma** = mid-anterior CC, and **Ca** = anterior CC. The lower panel displays the cerebellum segmentation based on the CERES atlas.

### 2.5 Segmentation of Cerebellar White and Grey Matter

For further parcellation of the cerebellum, a multi-atlas segmentation tool, CEREbellum Segmentation (CERES), was used to automatically parcellate the cerebellum lobule (Romero *et al*., 2017). CERES employs an optimised patch-based algorithm and demonstrates superior accuracy and execution time when parcellating the cerebellum compared to other pipelines (Romero *et al*., 2017). CERES includes the white and grey matter volumes of 12 cerebellar lobules, along with cortical thickness measurements (Fig. 1).

### 2.6 Statistical Analysis

Regional volume was selected as the primary measure for assessing structural changes. For within and between-subject comparisons, the volumetric regional measures were normalised by the Total Intracranial Volume (TIV) (Whitwell *et al*., 2001; Haar *et al*., 2016).

To assess volumetric changes in PD patients and healthy controls over time, we utilised Shapiro-Wilk tests to confirm non-normal data distribution, leading to the application of non-parametric Wilcoxon Signed-Rank and Mann-Whitney U tests for within-group and between-group comparisons, respectively. Python and Prism facilitated these analyses, with False Discovery Rate (FDR) correction used for multiple comparisons.

## 3. Results

### 3.1 Baseline Comparison

The region-specific volumetric comparisons in this study were done after normalising for TIV. TIV itself showed no significant differences between PD patients and controls, in the non-parametric Mann-Whitney U test (z = 0.10, p = 0.92), used due to non-normal distribution after testing the normality using Shapiro-Wilk test (Fig. 2). Subsequent analysis of specific ROIs normalised against TIV revealed significant volumetric differences in the bilateral pallidum, thalamus, and cerebellar white matter (q(FDR corrected significance) < 0.05), all showing higher volume in PD (Fig. 2). In the cerebellum and thalamus, the left hemisphere had a greater difference than the right. Anterior CC also depicted a marginal difference, whereby the patient group was higher than the control group (p = 0.02) though this significance did not survive FDR correction.

**Figure 2.**
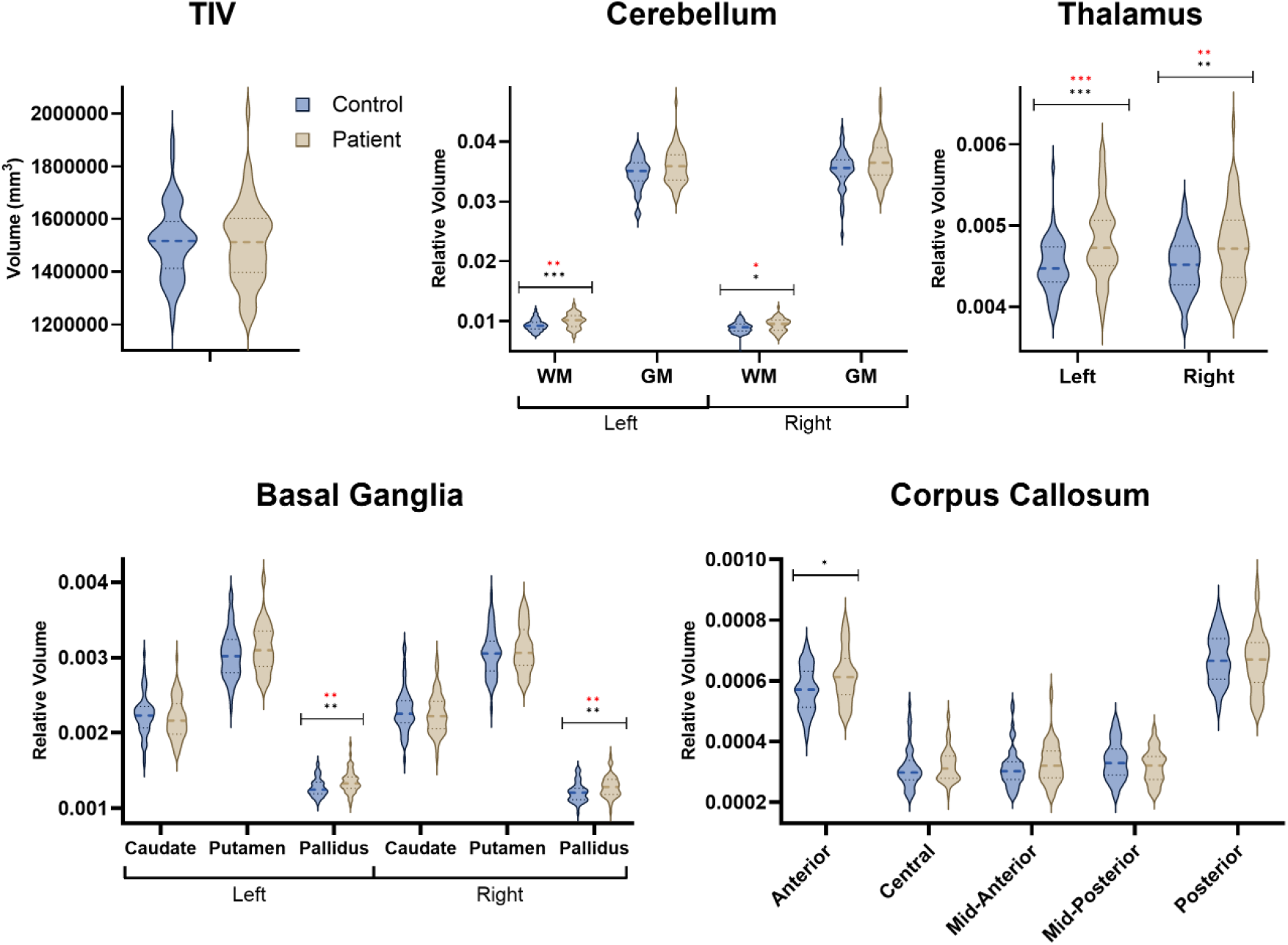
Baseline Comparison of TIV and Relative Volume in the Regions of Interest. The regional volume is presented as a portion of the TIV. Between-group differences were tested using the Mann-Whitney U test and FDR multiple comparisons correction. Significance levels are plotted by uncorrected p-values (black asterisk) and FDR-corrected q-values (red asterisk). *: values<0.05, **: values<0.01, ***: values<0.001. Cerebellum is labelled as: WM = white matter, GM = grey matter.

### 3.2 Longitudinal Volumetric Changes After 2/3 Years

To examine longitudinal changes over 2 to 3 years, we performed a one-sided Wilcoxon signed-rank test. The control group displayed a significant decline in TIV (q = 0.013) but not the PD group (q = 0.56). While TIV decline showed no significant difference between PD patients and controls (z = -2.23, q = 0.074), there was a trend of less severe decline in PD. Specific brain regions (Fig. 3 and Table S2) highlighted differential patterns of volumetric change (normalised for TIV). In healthy controls, notable decreases were observed in the thalamus (q= 0.007) and mid-CC (q < 0.001), suggesting these areas are particularly susceptible to changes associated with normal ageing. Conversely, the PD group demonstrated a volumetric decrease in the putamen (p <0.05, q = 0.06) and a most significant decrease in the cerebellar white matter (p = 0.002, q = 0.03). Comparative analyses between groups were done using the Mann-Whitney U-test. Multiple regions reported uncorrected significant differences including the right striatum and cerebellar white matter (p <0.05), but the only ROI showing significant group difference after correcting for multiple comparisons was the mid-posterior CC (q = 0.007).

**Figure 3.**
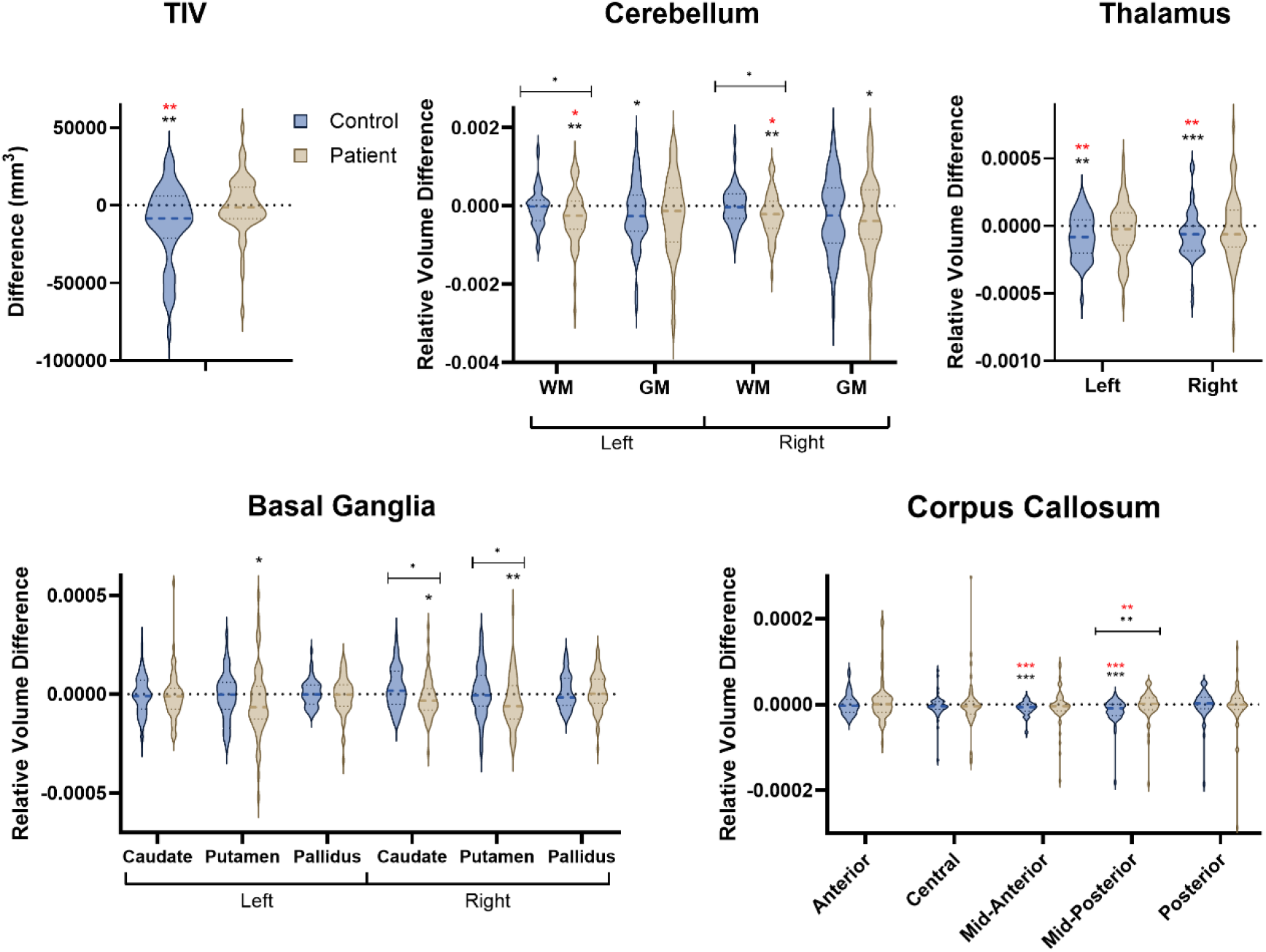
Change in the TIV and Relative Volume of the Selected Regions after 2/3 Years. The regional volume is presented as a portion of the TIV. Between-group differences were tested using the Mann-Whitney U test and FDR multiple comparisons correction. Significance levels are illustrated by uncorrected p-value (black asterisk) and corrected q-value (red asterisk). *: values<0.05, **: values<0.01, ***: values<0.001. Cerebellum is labelled as: WM = white matter, GM = grey matter.

### 3.3 Relative Volume Comparison in the Cerebellum in PD

To better assess group differences in the observed cerebellar decline, we used the CERES pipeline to parcellate the cerebellar volume into 12 lobules. The white matter in Crus I and lobule IX showed the most significant decline, with Crus I showing a notable volume decrease in both cerebellar hemispheres (left: q = 0.02, right: q < 0.001) and lobule IX displaying significant decline in the right hemisphere (q = 0.02) while the decline in left lobule IX did not survive FDR correction (p<0.02, q=0.088). Additional cerebellar regions exhibited marginal significance that did not withstand rigorous multiple comparison corrections (Fig. 4 and Tables S3 and S4).

**Figure 4.**
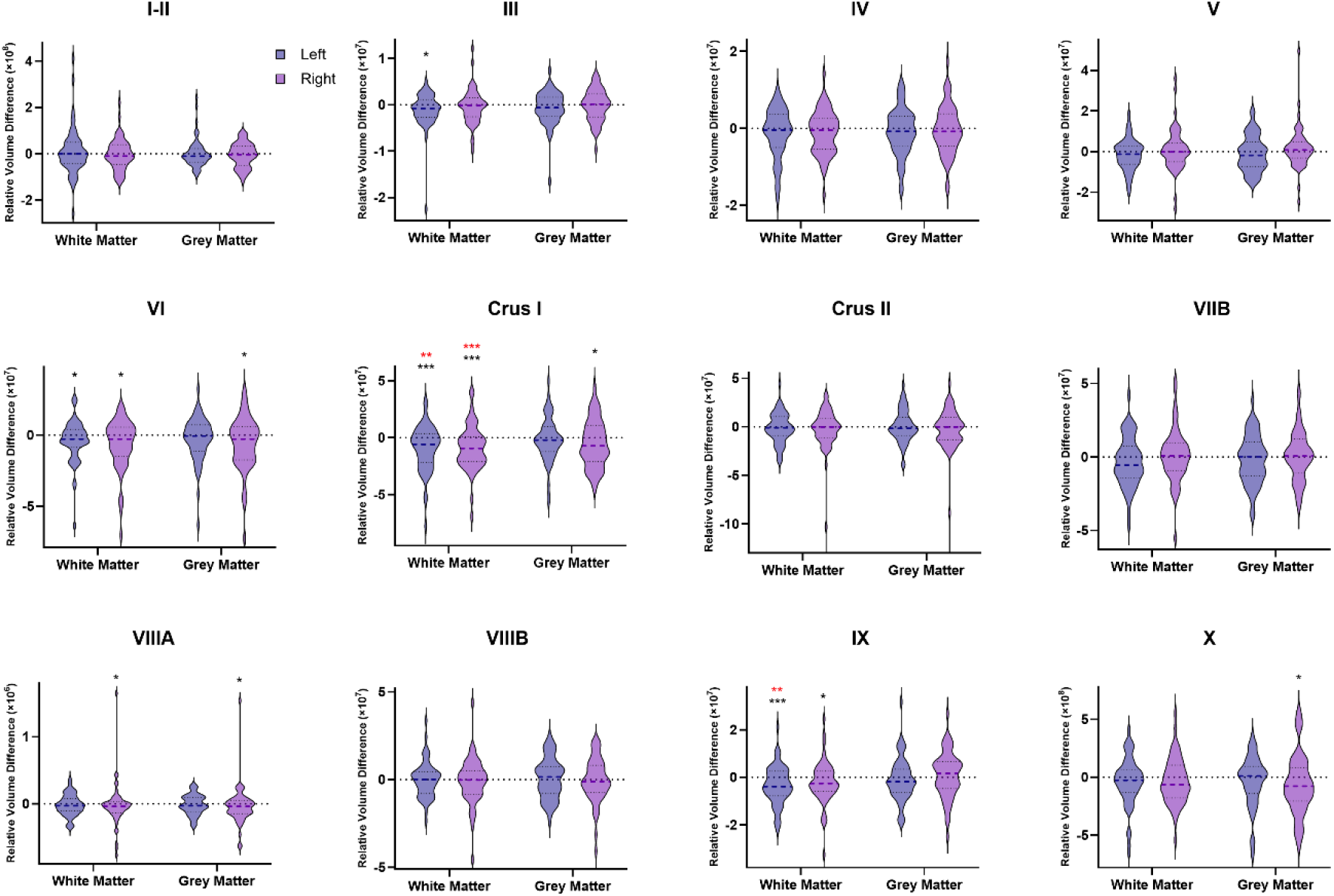
Change in Relative Volume in the Cerebellum of PD Patient after 2/3 Years. The regional volume was normalised to the TIV. Within-group differences were tested using the Wilcoxon test and FDR multiple comparisons. Significance levels were represented by uncorrected p-values (black asterisk) and corrected q-values (red asterisk). *: values<0.05, **: values<0.01, ***: values<0.001.

## 4. Discussion

In this study, we explored volumetric changes in subcortical grey and white matter across a 2 to 3-year interval in early PD patients alongside an age-matched healthy control cohort. The longitudinal approach and the inclusion of healthy controls, matched for confounding factors, allowed a focused longitudinal examination of regional volumetric alterations. Our findings underline the crucial role of the cerebellum in early PD progression. We showed that during baseline, shortly after PD diagnosis, patients had increased volume in cerebellar white matter, but over the following 2-3 years, this area showed the most significant decrease in volume. These results suggest a potential compensatory role of the cerebellum in prodromal and early PD which changes around diagnostic and symptoms appearance. We further delved into cerebellar lobule segmentation and localised this volume decrease in Crus I, which was previously suggested to be a hub of the functional cerebellar connectome (Chen *et al*., 2022).

While there were no significant differences in the Total Intracranial Volume between the early PD patients and matching controls at baseline, PD patients displayed a higher volume in multiple subcortical regions, including the cerebellar white matter, pallidum, and thalamus (Whone *et al*., 2003). This may indicate the initial compensatory mechanism in prodromal and early PD (Brotchie and Fitzer-Attas, 2009; Blesa *et al*., 2017). A few previous studies reported differences in pallidum volume between control and PD groups, though with no statistical significance (Yang *et al*., 2024). While some previous studies found no detectable regional atrophy differences in the thalamus of PD patients and controls (McKeown *et al*., 2008; Ibarretxe-Bilbao *et al*., 2012; ‘Progression of brain atrophy in the early stages of Parkinson’s disease: A longitudinal tensor-based morphometry study in de novo patients without cognitive impairment’, no date), others reported an increased thalamic volume in early PD patients (Jia *et al*., 2015). A more recent study looking into all volume, thickness, and surface area of the thalamus highlighted an overall bigger thalamus in early PD patients (Kerestes *et al*., 2023). The significantly higher cerebellar white matter volume in PD patients is consistent with a recent observation of a higher anterior cerebellar volume in the initial stage of PD compared to controls. It is also in line with the proposed hyperactivation of the cerebellothalamic pathway explained by basal ganglia dysfunction (Helmich *et al*., 2011). Recent studies demonstrate a growing interest in understanding the reciprocal interaction between the cerebellum and basal ganglia (Bostan and Strick, 2018). An anatomical tracing study underlined both cerebellum-striatum and subthalamic nucleus-cerebellum networks via the thalamus and pontine nucleus, respectively (Washburn *et al*., 2024). Our findings in increased cerebellar white matter volume in early PD patients may suggest basal ganglia-mediated hyperactivation, which is shown to be relevant in understanding PD tremors as well (Helmich *et al*., 2011).

Our longitudinal analysis over a 2–3-year period suggests that the TIV decreases as part of healthy ageing, though this decay was significant only within the control group, but not in the PD group. Yet, the difference in the decay between the groups lacked significance. This is consistent with earlier findings of increased TIV in PD (Krabbe *et al*., 2005). In the studied ROIs, the overall trend was of volumetric decay but while some regions showed more severe decay in PD others showed more severe decay in the control group. Healthy controls demonstrated the greatest reduction in the mid-CC and thalamus, whereas the PD patients had the biggest reduction in the volume of the cerebellar white matter. The mid-posterior CC was identified as a region with the greatest difference between the groups, showing greater decay in healthy controls. A recent longitudinal study in PD reported no structural change in the CC in a 21-month follow-up (Wu *et al*., 2020); we also found no change in PD but show that this is in contrast to a significant change in healthy ageing. This highlights the importance of healthy control comparison in longitudinal patient studies.

In PD patients, the biggest change was observed in the cerebellar white matter, which is consistent with the pathological α-synuclein aggregates found in the lobular white matter tracts (Seidel *et al*., 2017). One of the main hypotheses revolving around the selective neuronal vulnerability observed in PD is based on α-synuclein pathology, where insoluble synuclein aggregation can disseminate along neural pathways in a prion-like mechanism. Previously, studies have shown that Lewy bodies formed in the Substantia Nigra positively correlates with the symptom severity (Hartmann, 2004). The presence of α-synuclein aggregates in the cerebellum along with our findings suggest an importance in investigating early pathology in the cerebellum.

Kerestes et al., (2023) also observed notable changes in the cerebellar volume based on HY score increment. Interestingly, the cerebellum is considered to be one of the brain areas which undergoes the slowest neurodegeneration (Liang and Carlson, 2020) and to play a role in compensatory mechanism in early PD (Simioni, Dagher and Fellows, 2016). Its fast WM volumetric decay presented here might capture the transition from its compensatory role to its neurodegeneration towards distinct cerebellar atrophy In advanced PD (O’Callaghan *et al*., 2016).

Looking deeper into the cerebellar volumetric decay, our results suggest that bilateral Crus I and the left IX lobule have the greatest volumetric loss in PD after a 2/3-year follow-up. Multiple studies observed degeneration in Crus I in advanced PD patients (O’Callaghan *et al*., 2016; Erro *et al*., 2020) and one classified PD patients from matched controls based on cerebellum Crus I grey matter degeneration with 95% accuracy (Zeng *et al*., 2017). Ongoing research continues to uncover cerebellar macrostructural changes in PD relative to cognitive impairments. A recent study using the same segmentation tool as ours, CERES, found a strong association between the cerebellum and cognitive function (Bègue *et al*., 2023). In particular, Crus I showed a positive association with working memory. Hence, the decreased Crus I WM volume in our findings may explain the developing non-motor symptoms in PD patients. Moreover, enhanced Crus I activity and its increased connectivity with the putamen are positively correlated with cognitive assessments like MoCA, indicating the region’s role in cognitive impairments in PD (Shen *et al*., 2020; Solstrand Dahlberg, Lungu and Doyon, 2020). This is further supported by the recent findings by Kerestes et al. (2023), which underlined the association between cognitive performance and the decline in cerebellar volume, suggesting the critical link between the cerebellum and cognitive function in PD.

There are limited reported findings on lobule IX in PD pathology, though one study observed a difference in lobule IX between PD and healthy controls (Gardoni *et al*., 2023). The change in lobule IX does not seem to have a strict relationship with either motor or non-motor symptomatology. Rather, the current literature suggests a more intricate participation in both. For instance, one study found a strong correlation between depression, anxiety, and IX atrophy (Ma *et al*., 2018). This is further explained by the involvement of the cerebellum in the cortico-limbic network, regulating the negative emotion processing (Adamaszek *et al*., 2017). Some also suggest an involvement of lobule IX in visuospatial impairment in PD (Lawn and ffytche, 2021; Yin *et al*., 2021). In contrast, a few studies suggest an increased IX activity in dopamine-resistant PD tremor (Dirkx *et al*., 2019). This is particularly relevant as the cerebellum consists mainly of non-dopaminergic transmitters (Zhong *et al*., 2022). Similarly, Lopez and colleagues emphasised the relevance of XI in predicting resting tremors in PD patients (Lopez *et al*., 2020). Nonetheless, a further investigation of the involvement of lobule IX in PD symptomology is required.

While our study presents the most complete analysis of anatomical changes in PD to date, combining cross-sectional and longitudinal design, it was subject to several known limitations. First, our results are based on Freesurfer segmentation, which while considered state-of-the-art, when comparing pathological conditions with controls some issues were found in some of its previous versions (Filip *et al*., 2022). Our study implemented the recently updated Freesurfer version 7.3.2, which should be more robust, but its replicability and reproducibility have not yet been externally tested and reported. Second, the MRI data for the PD patients and controls were obtained from different multisite datasets. Though we controlled for TE and TR variability, there are likely additional confounding factors. There was also high variability in voxel dimensions within the same dataset that needed to be controlled for. Thirdly, the control imaging data acquired from OASIS-3 were not fully compatible with the automated CERES pipeline. As we were not authorised to modify the commands used in the CERES pipeline, the cerebellar segmentation for controls was not included in our analysis. However, our initial subcortical segmentation using Freesurfer demonstrated overall insignificant changes in the cerebellum white matter. This is also consistent with an epigenetic study, where the ageing in the cerebellum is slower than in other body parts (Horvath *et al*., 2015).

In conclusion, anatomical MRI data of PD patients and matching controls at baseline and 2/3-year follow-up, demonstrate significantly different volumetric decay between the groups. Our approach revealed that the cerebellar WM showed significantly increased atrophy in PD relative to healthy ageing. This was driven by atrophy in cerebellar Crus I and IX white matter in early PD pathology. Furthermore, our analysis enabled us to explore potential compensatory increases in volume in the thalamus and mid-CC in early PD, relative to changes observed in healthy ageing. Future data collection efforts over longer periods with shorter intervals will enhance our understanding of structural changes during PD progression. Future work should aim to comprehensively identify morphological brain changes that occur before the onset of PD symptoms to improve the understanding of PD pathology.

## Acknowledgements

The study was supported by the UK Dementia Research Institute, Care Research & Technology Centre and SH’s Edmond and Lily Safra Fellowship. The PD data used in the preparation of this article were obtained from the Parkinson’s Progression Markers Initiative (PPMI) database (https://www.ppmi-info.org). PPMI – a public-private partnership – is funded by The Michael J. Fox Foundation for Parkinson’s Research and funding partners listed here https://www.ppmi-info.org/about-ppmi/who-we-are/studysponsors. The HC data were obtained from the Open Access Series of Imaging Series 3 (OASIS-3, https://sites.wustl.edu/oasisbrains/home/oasis-3).

## Supplementary Materials

**Table S1.** MRI Scanning Parameters of PPMI and OASIS-3.

|  | PPMI | OASIS-3 | T-statistic | p-value |
| --- | --- | --- | --- | --- |
| Field Strength | 3.0 | 3.0 | - | - |
| Manufacturer | Siemens | Siemens | - | - |
| Echo Time (ms) | 3.00 ±0.02 | 3.39 ±3.7 | 1.11 | 0.27 |
| Repetition Time (ms) | 2280 ±217 | 2437 ±839 | 1.93 | 0.054 |

**Table S2.**
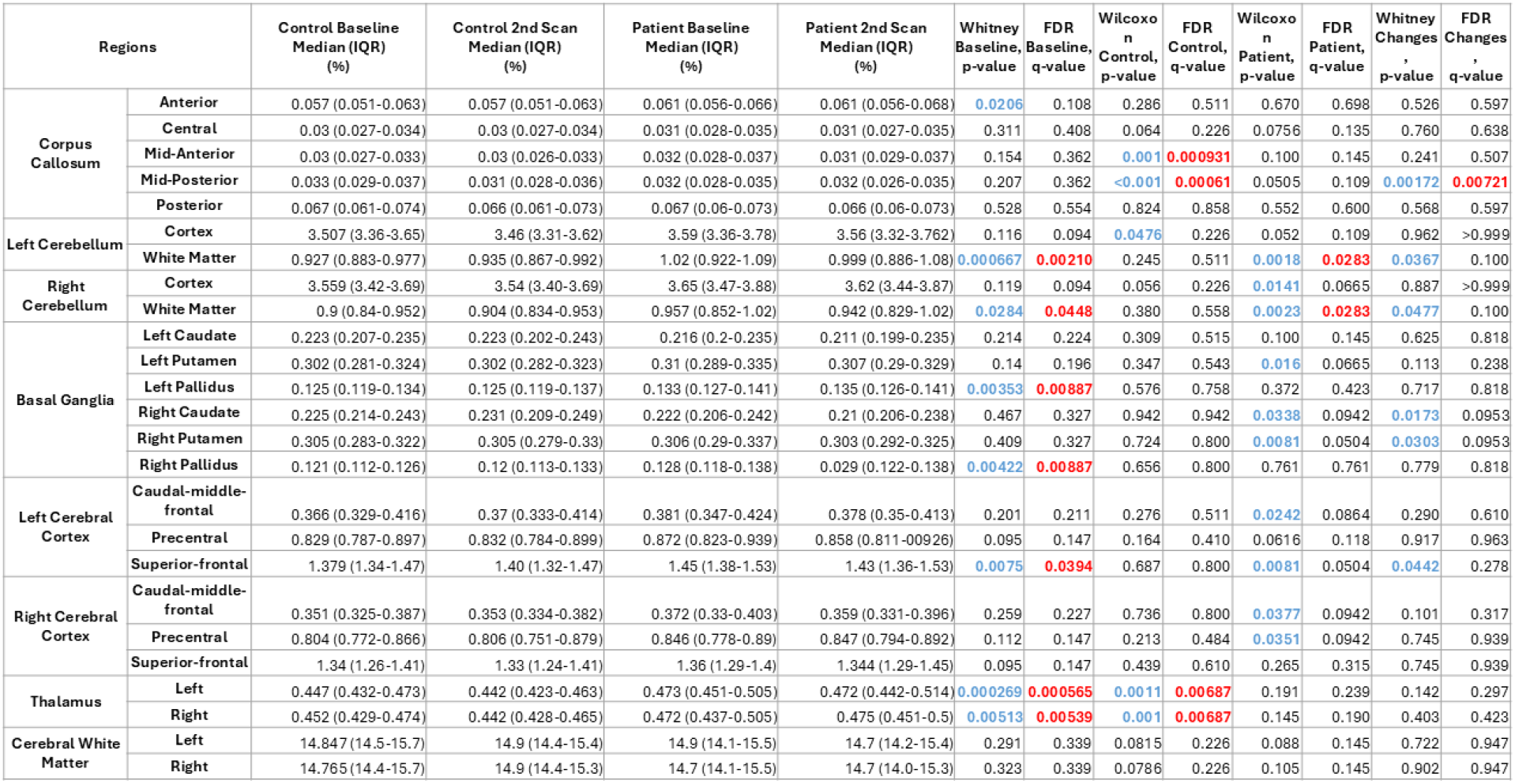
Baseline and Relative Volumetric Changes. Full volumetric data (median and IQR for each ROI) for both groups in both scans, including the control baseline median with inter-quartile range (IQR), control 2nd scan median (IQR), patient baseline median (IQR), patient 2nd scan median (IQR), U-test baseline control vs. patient (p-value), Wilcoxon test control (p-value), Wilcoxon test patient (p-value), U-test changes control vs. patient (p-value), and their respective FDR-corrected q-values. Values highlighted by blue indicate significant uncorrected p-values, whereas red represents significant corrected q-values. The relative median and its interquartile range (IQR) values were converted into percentages for easier reading.

**Table S3.** Relative Volumetric Changes in the Cerebellar White Matter. The baseline median (IQR), 2nd scan median (IQR), Wilcoxon p-value, and the FDR-corrected q-value for the left and right of each region. Values in blue represent significant uncorrected p-values, whereas red indicates significant corrected q-values. The relative median and its IQR values were multiplied by 1e5 for the simplicity of reading.

| Regions |  | Baseline Median (IQR) (x1e5) | 2nd Scan Median (IQR) (x1e5) | Wilcoxon, p-value | FDR, q-value |
| --- | --- | --- | --- | --- | --- |
| I-II | Left | 0.004 (0.003-0.004) | 0.004 (0.003-0.004) | 0.513 | 0.597 |
|  | Right | 0.004 (0.004-0.005) | 0.004 (0.003-0.005) | 0.247 | 0.358 |
| III | Left | 0.047 (0.04-0.054) | 0.046 (0.041-0.052) | **0.011** | 0.088 |
|  | Right | 0.049 (0.043-0.055) | 0.048 (0.043-0.054) | 0.146 | 0.292 |
| IV | Left | 0.16 (0.143-0.176) | 0.16 (0.144-0.175) | 0.203 | 0.35 |
|  | Right | 0.151 (0.143-0.166) | 0.151 (0.141-0.165) | 0.108 | 0.292 |
| V | Left | 0.275 (0.254-0.3) | 0.271 (0.253-0.301) | 0.063 | 0.21 |
|  | Right | 0.26 (0.248-0.296) | 0.266 (0.245-0.292) | 0.59 | 0.641 |
| VI | Left | 0.547 (0.527-0.611) | 0.542 (0.518-0.594) | **0.033** | 0.127 |
|  | Right | 0.557 (0.511-0.627) | 0.553 (0.506-0.606) | **0.013** | 0.088 |
| Crus I | Left | 0.84 (0.746-0.923) | 0.811 (0.73-0.924) | **0.001** | **0.017** |
|  | Right | 0.852 (0.788-0.946) | 0.836 (0.782-0.944) | **<0.001** | **<0.001** |
| Crus II | Left | 0.549 (0.469-0.642) | 0.544 (0.474-0.642) | 0.254 | 0.358 |
|  | Right | 0.567 (0.517-0.644) | 0.569 (0.509-0.631) | 0.265 | 0.358 |
| VIIB | Left | 0.314 (0.287-0.35) | 0.308 (0.283-0.353) | 0.061 | 0.21 |
|  | Right | 0.323 (0.296-0.352) | 0.316 (0.298-0.356) | 0.612 | 0.651 |
| VIII A | Left | 0.4 (0.351-0.458) | 0.398 (0.357-0.449) | 0.141 | 0.292 |
|  | Right | 0.393 (0.358-0.416) | 0.383 (0.347-0.416) | **0.023** | 0.104 |
| VIII B | Left | 0.271 (0.247-0.303) | 0.27 (0.244-0.307) | 0.41 | 0.512 |
|  | Right | 0.264 (0.238-0.299) | 0.264 (0.237-0.297) | 0.369 | 0.473 |
| IX | Left | 0.242 (0.214-0.272) | 0.237 (0.212-0.271) | **0.001** | **0.017** |
|  | Right | 0.249 (0.228-0.275) | 0.247 (0.223-0.276) | **0.013** | 0.088 |
| X | Left | 0.042 (0.037-0.044) | 0.041 (0.038-0.043) | 0.143 | 0.292 |
|  | Right | 0.043 (0.039-0.045) | 0.042 (0.039-0.045) | 0.072 | 0.225 |

**Table S4.** Relative Volumetric Changes in the Cerebellar Grey Matter. The baseline median (IQR), 2nd scan median (IQR), Wilcoxon p-value, and the FDR-corrected q-value for the left and right of each region. Values in blue indicate significant uncorrected p-values, whereas red represents significant corrected q-values. The relative median and their IQR values were multiplied by 1e5 for the simplicity of reading.

| Regions |  | Baseline Median (IQR) (x1e5) | 2nd Scan Median (IQR) (x1e5) | Wilcoxon, p-value | FDR, q-value |
| --- | --- | --- | --- | --- | --- |
| I-II | Left | 0.002 (0.001-0.002) | 0.002 (0.001-0.002) | 0.135 | 0.292 |
|  | Right | 0.002 (0.002-0.003) | 0.002 (0.002-0.003) | 0.236 | 0.358 |
| III | Left | 0.034 (0.029-0.038) | 0.033 (0.027-0.038) | 0.117 | 0.292 |
|  | Right | 0.033 (0.028-0.038) | 0.034 (0.029-0.038) | 0.5 | 0.595 |
| IV | Left | 0.141 (0.127-0.156) | 0.139 (0.127-0.154) | 0.162 | 0.312 |
|  | Right | 0.128 (0.122-0.144) | 0.131 (0.119-0.143) | 0.265 | 0.358 |
| V | Left | 0.243 (0.224-0.266) | 0.238 (0.225-0.261) | 0.144 | 0.292 |
|  | Right | 0.222 (0.206-0.245) | 0.225 (0.206-0.246) | 0.854 | 0.854 |
| VI | Left | 0.495 (0.481-0.556) | 0.501 (0.477-0.543) | 0.176 | 0.314 |
|  | Right | 0.495 (0.445-0.558) | 0.495 (0.446-0.542) | 0.022 | 0.104 |
| Crus I | Left | 0.706 (0.639-0.792) | 0.701 (0.634-0.798) | 0.17 | 0.314 |
|  | Right | 0.718 (0.652-0.79) | 0.708 (0.636-0.787) | 0.025 | 0.104 |
| Crus II | Left | 0.471 (0.41-0.556) | 0.474 (0.41-0.562) | 0.526 | 0.598 |
|  | Right | 0.49 (0.442-0.55) | 0.489 (0.431-0.547) | 0.231 | 0.358 |
| VII B | Left | 0.281 (0.252-0.306) | 0.272 (0.243-0.309) | 0.254 | 0.358 |
|  | Right | 0.282 (0.264-0.313) | 0.282 (0.261-0.316) | 0.574 | 0.638 |
| VIII A | Left | 0.354 (0.313-0.394) | 0.353 (0.312-0.391) | 0.224 | 0.358 |
|  | Right | 0.344 (0.31-0.366) | 0.338 (0.306-0.362) | 0.011 | 0.088 |
| VIII B | Left | 0.231 (0.208-0.258) | 0.23 (0.209-0.262) | 0.673 | 0.701 |
|  | Right | 0.231 (0.204-0.257) | 0.227 (0.206-0.256) | 0.451 | 0.55 |
| IX | Left | 0.195 (0.171-0.21) | 0.192 (0.17-0.218) | 0.135 | 0.292 |
|  | Right | 0.205 (0.183-0.222) | 0.2 (0.187-0.224) | 0.793 | 0.809 |
| X | Left | 0.039 (0.035-0.042) | 0.039 (0.035-0.041) | 0.284 | 0.374 |
|  | Right | 0.039 (0.035-0.041) | 0.037 (0.034-0.041) | 0.014 | 0.088 |

## References

Abulikemu, S., Tai, Y.F. and Haar, S. (2023) ‘Modulatory Effect of Levodopa on the Basal Ganglia-Cerebellum Connectivity in Parkinson’s Disease’. bioRxiv, P. 2023.01.16.524229. Available at: 10.1101/2023.01.16.524229.

Adamaszek, M. et al. (2017) ‘Consensus Paper: Cerebellum and Emotion’, The Cerebellum, 16(2), pp. 552–576. Available at: 10.1007/s12311-016-0815-8.

Bègue, I. et al. (2023) ‘The Cerebellum and Cognitive Function: Anatomical Evidence from a Transdiagnostic Sample’, The Cerebellum [Preprint]. Available at: 10.1007/s12311-023-01645-y.

Blesa, J. et al. (2017) ‘Compensatory mechanisms in Parkinson’s disease: Circuits adaptations and role in disease modification’, Experimental Neurology, 298, pp. 148–161. Available at: 10.1016/j.expneurol.2017.10.002.

Bostan, A.C. and Strick, P.L. (2018) ‘The basal ganglia and the cerebellum: nodes in an integrated network’, Nature Reviews Neuroscience, 19(6), pp. 338–350. Available at: 10.1038/s41583-018-0002-7.

Brotchie, J. and Fitzer-Attas, C. (2009) ‘Mechanisms compensating for dopamine loss in early Parkinson disease’, Neurology, 72(7_supplement_2). Available at: 10.1212/WNL.0b013e318198e0e9.

Chen, Z. et al. (2022) ‘Functional connectome of human cerebellum’, NeuroImage, 251, p. 119015. Available at: 10.1016/j.neuroimage.2022.119015.

Dirkx, M.F. et al. (2019) ‘Cerebral differences between dopamine-resistant and dopamine-responsive Parkinson’s tremor’, Brain, 142(10), pp. 3144–3157. Available at: 10.1093/brain/awz261.

Erro, R. et al. (2020) ‘Subcortical atrophy and perfusion patterns in Parkinson disease and multiple system atrophy’, Parkinsonism & Related Disorders, 72, pp. 49–55. Available at: 10.1016/j.parkreldis.2020.02.009.

Filip, P. et al. (2022) ‘Different FreeSurfer versions might generate different statistical outcomes in case–control comparison studies’, Neuroradiology, 64(4), pp. 765–773. Available at: 10.1007/s00234-021-02862-0.

Fischl, B. (2012) ‘FreeSurfer’, NeuroImage, 62(2), pp. 774–781. Available at: 10.1016/j.neuroimage.2012.01.021.

Gardoni, A. et al. (2023) ‘Cerebellar alterations in Parkinson’s disease with postural instability and gait disorders’, Journal of Neurology, 270(3), pp. 1735–1744. Available at: 10.1007/s00415-022-11531-y.

Haar, S. et al. (2016) ‘Anatomical Abnormalities in Autism?’, Cerebral Cortex, 26(4), pp. 1440–1452. Available at: 10.1093/cercor/bhu242.

Hartmann, A. (2004) ‘Postmortem studies in Parkinson’s disease’, Dialogues in Clinical Neuroscience, 6(3), pp. 281–293.

Helmich, R.C. et al. (2011) ‘Pallidal dysfunction drives a cerebellothalamic circuit into Parkinson tremor’, Annals of Neurology, 69(2), pp. 269–281. Available at: 10.1002/ana.22361.

Horvath, S. et al. (2015) ‘The cerebellum ages slowly according to the epigenetic clock’, Aging, 7(5), pp. 294–306. Available at: 10.18632/aging.100742.

Ibarretxe-Bilbao, N. et al. (2012) ‘Progression of cortical thinning in early Parkinson’s disease’, Movement Disorders, 27(14), pp. 1746–1753. Available at: 10.1002/mds.25240.

Jia, X. et al. (2015) ‘Longitudinal Study of Gray Matter Changes in Parkinson Disease’, AJNR: American Journal of Neuroradiology, 36(12), pp. 2219–2226. Available at: 10.3174/ajnr.A4447.

Johansson, M.E. et al. (2024) ‘Clinical severity in Parkinson’s disease is determined by decline in cortical compensation’, Brain, 147(3), pp. 871–886. Available at: 10.1093/brain/awad325.

Kerestes, R. et al. (2023) ‘Cerebellar Volume and Disease Staging in Parkinson’s Disease: An ENIGMA-PD Study’, Movement Disorders, 38(12), pp. 2269–2281. Available at: 10.1002/mds.29611.

Krabbe, K. et al. (2005) ‘Increased intracranial volume in Parkinson’s disease’, Journal of the Neurological Sciences, 239(1), pp. 45–52. Available at: 10.1016/j.jns.2005.07.013.

Lawn, T. and ffytche, D. (2021) ‘Cerebellar correlates of visual hallucinations in Parkinson’s disease and Charles Bonnet Syndrome’, Cortex, 135, pp. 311–325. Available at: 10.1016/j.cortex.2020.10.024.

Li, T., Le, W. and Jankovic, J. (2023) ‘Linking the cerebellum to Parkinson disease: an update’, Nature Reviews Neurology, 19(11), pp. 645–654. Available at: 10.1038/s41582-023-00874-3.

Liang, K.J. and Carlson, E.S. (2020) ‘Resistance, vulnerability and resilience: A review of the cognitive cerebellum in aging and neurodegenerative diseases’, Neurobiology of Learning and Memory, 170, p. 106981. Available at: 10.1016/j.nlm.2019.01.004.

Lopez, A.M. et al. (2020) ‘Structural Correlates of the Sensorimotor Cerebellum in Parkinson’s Disease and Essential Tremor’, Movement Disorders, 35(7), pp. 1181–1188. Available at: 10.1002/mds.28044.

Ma, X. et al. (2018) ‘Cerebellar atrophy in different subtypes of Parkinson’s disease’, Journal of the Neurological Sciences, 392, pp. 105–112. Available at: 10.1016/j.jns.2018.06.027.

Marsden, C.D. (1990) ‘Parkinson’s disease’, Lancet (London, England), 335(8695), pp. 948–952. Available at: 10.1016/0140-6736(90)91006-v.

McKeown, M.J. et al. (2008) ‘Shape (but not volume) changes in the thalami in Parkinson disease’, BMC Neurology, 8, p. 8. Available at: 10.1186/1471-2377-8-8.

Nazem, S. et al. (2009) ‘Montreal Cognitive Assessment Performance in Patients with Parkinson’s Disease with “Normal” Global Cognition According to Mini-Mental State Examination Score’, Journal of the American Geriatrics Society, 57(2), pp. 304–308. Available at: 10.1111/j.1532-5415.2008.02096.x.

O’Callaghan, C. et al. (2016) ‘Cerebellar atrophy in Parkinson’s disease and its implication for network connectivity’, Brain, 139(3), pp. 845–855. Available at: 10.1093/brain/awv399.

Pitcher, T.L. et al. (2012) ‘Reduced striatal volumes in Parkinson’s disease: a magnetic resonance imaging study’, Translational Neurodegeneration, 1(1), p. 17. Available at: 10.1186/2047-9158-1-17.

‘Progression of brain atrophy in the early stages of Parkinson’s disease: A longitudinal tensor-based morphometry study in de novo patients without cognitive impairment’ (no date). Available at: 10.1002/hbm.22449.

Romero, J.E. et al. (2017) ‘CERES: A new cerebellum lobule segmentation method’, NeuroImage, 147, pp. 916–924. Available at: 10.1016/j.neuroimage.2016.11.003.

Seidel, K. et al. (2017) ‘Involvement of the cerebellum in Parkinson disease and dementia with Lewy bodies’, Annals of Neurology, 81(6), pp. 898–903. Available at: 10.1002/ana.24937.

Shen, B. et al. (2020) ‘Altered putamen and cerebellum connectivity among different subtypes of Parkinson’s disease’, CNS Neuroscience & Therapeutics, 26(2), pp. 207–214. Available at: 10.1111/cns.13259.

Simioni, A.C., Dagher, A. and Fellows, L.K. (2016) ‘Compensatory striatal–cerebellar connectivity in mild–moderate Parkinson’s disease’, NeuroImage: Clinical, 10, pp. 54–62. Available at: 10.1016/j.nicl.2015.11.005.

Solstrand Dahlberg, L., Lungu, O. and Doyon, J. (2020) ‘Cerebellar Contribution to Motor and Non-motor Functions in Parkinson’s Disease: A Meta-Analysis of fMRI Findings’, Frontiers in Neurology, 11. Available at: 10.3389/fneur.2020.00127 (Accessed: 15 August 2023).

Szewczyk-Krolikowski, K. et al. (2014) ‘Functional connectivity in the basal ganglia network differentiates PD patients from controls’, Neurology, 83(3), p. 208. Available at: 10.1212/WNL.0000000000000592.

Taylor, K.I. et al. (2018) ‘Progressive Decline in Gray and White Matter Integrity in de novo Parkinson’s Disease: An Analysis of Longitudinal Parkinson Progression Markers Initiative Diffusion Tensor Imaging Data’, Frontiers in Aging Neuroscience, 10. Available at: 10.3389/fnagi.2018.00318 (Accessed: 15 August 2023).

Washburn, S. et al. (2024) ‘The cerebellum directly modulates the substantia nigra dopaminergic activity’, Nature Neuroscience, 27(3), pp. 497–513. Available at: 10.1038/s41593-023-01560-9.

Whitwell, J.L. et al. (2001) ‘Normalization of Cerebral Volumes by Use of Intracranial Volume: Implications for Longitudinal Quantitative MR Imaging’, AJNR: American Journal of Neuroradiology, 22(8), pp. 1483–1489.

Whone, A.L. et al. (2003) ‘Plasticity of the nigropallidal pathway in Parkinson’s disease’, Annals of Neurology, 53(2), pp. 206–213. Available at: 10.1002/ana.10427.

Wu, J. et al. (2020) ‘Longitudinal Macro/Microstructural Alterations of Different Callosal Subsections in Parkinson’s Disease Using Connectivity-Based Parcellation’, Frontiers in Aging Neuroscience, 12. Available at: 10.3389/fnagi.2020.572086 (Accessed: 15 August 2023).

Yang, W. et al. (2024) ‘The longitudinal volumetric and shape changes of subcortical nuclei in Parkinson’s disease’, Scientific Reports, 14(1), p. 7494. Available at: 10.1038/s41598-024-58187-4.

Yin, K. et al. (2021) ‘Resting-state functional magnetic resonance imaging of the cerebellar vermis in patients with Parkinson’s disease and visuospatial disorder’, Neuroscience Letters, 760, p. 136082. Available at: 10.1016/j.neulet.2021.136082.

Zeng, L.-L. et al. (2017) ‘Differentiating Patients with Parkinson’s Disease from Normal Controls Using Gray Matter in the Cerebellum’, The Cerebellum, 16(1), pp. 151–157. Available at: 10.1007/s12311-016-0781-1.

Zhong, Y. et al. (2022) ‘A review on pathology, mechanism, and therapy for cerebellum and tremor in Parkinson’s disease’, npj Parkinson’s Disease, 8(1), pp. 1–9. Available at: 10.1038/s41531-022-00347-2.

